# A framework for research into continental ancestry groups of the UK Biobank

**DOI:** 10.1101/2021.12.14.472589

**Authors:** Andrei-Emil Constantinescu, Ruth E. Mitchell, Jie Zheng, Caroline J. Bull, Nicholas J. Timpson, Borko Amulic, Emma E. Vincent, David A. Hughes

## Abstract

**Background:** The UK Biobank is a large prospective cohort, based in the United Kingdom, that has deep phenotypic and genomic data on roughly a half a million individuals. Included in this resource are data on approximately 78,000 individuals with “non-white British ancestry”. Whilst most epidemiology studies have focused predominantly on populations of European ancestry, there is an opportunity to contribute to the study of health and disease for a broader segment of the population by making use of the UK Biobank’s “non-white British ancestry” samples. Here we present an empirical description of the continental ancestry and population structure among the individuals in this UK Biobank subset.

**Results:** Reference populations from the 1000 Genomes Project for Africa, Europe, East Asia, and South Asia were used to estimate ancestry for each individual. Those with at least 80% ancestry in one of these four continental ancestry groups were taken forward (N=62,484). Principal component and K-means clustering analyses were used to identify and characterize population structure within each ancestry group. Of the approximately 78,000 individuals in the UK Biobank that are of “non-white British” ancestry, 50,685, 6,653, 2,782, and 2,364 individuals were associated to the European, African, South Asian, and East Asian continental ancestry groups, respectively. Each continental ancestry group exhibits prominent population structure that is consistent with self-reported country of birth data and geography.

**Conclusions:** Methods outlined here provide an avenue to leverage UK Biobank’s deeply phenotyped data allowing researchers to maximise its potential in the study of health and disease in individuals of non-white British ancestry.

## Introduction

As the research community strives to understand the genetic architecture of disease [1], it has increasingly realized the necessity of inclusion and diversity – of ethnically, ancestrally, environmentally, and geographically diverse populations [2–5]. Not simply to enhance knowledge about health and disease, but to insure health equity. Epidemiological studies, including genome-wide associations studies (GWAS), have been overwhelmingly conducted in European populations [2]. However, funding efforts and studies including the Human Heredity and Health in Africa (H3Africa) Initiative [6], the Population Architecture using Genomics and Epidemiology (PAGE) Consortium [7], Trans-Omics for Precision Medicine Consortium [8], Hispanic Community Health Study / Study of Latinos (SOL) [9], and the All of Us Research Program [10] are making concerted efforts to include and increase the number of under-represented populations in genomic epidemiology studies.

The UK Biobank project (UKBB) has phenotypic and genomic data from a prospective cohort of approximately 500,000 individuals from across the United Kingdom [11,12]. It has become an outstanding resource for studies of health and disease, and genetic diversity within the United Kingdom. Whilst it is made up of around 430,000 “white British ancestry” individuals, as defined by UKBB, it also contains a wealth of diversity from other self-described ethnicities (~78,000). This is a resource that should be utilized to help expand inclusion and diversity in epidemiological studies.

The Pan-UK Biobank, or the Pan-ancestry genetic analysis of the UKBB, has leveraged the diversity present in UKBB and is freely providing GWAS summary statistics for over seven thousand phenotypes in six continental ancestry groups (https://pan.ukbb.broadinstitute.org). The genetic “ancestry” groups identified by Pan-UK Biobank and within our study refer to groups of individuals with a shared genetic ancestry and demographic history. Studies and public resources like Pan-UK Biobank are vital to the goal of increasing under-represented populations and the larger goal of describing and understanding the genetic architecture of phenotypic traits and disease. However, the limited information on intra-population structure and non-specific use of covariates in Pan-UK Biobank GWAS models may influence association effect estimates. A description of the continental diversity and population structure present in the UKBB will aid future study design, methodological choice(s) and ultimately improve our understanding of how genotype influences phenotype.

Here, we describe an approach to define continental ancestry groups and provide a description of the structure and population differentiation within them. We define “ancestry” here as genetic ancestry or the complex inheritance of one’s genetic material, but in practice we will be using methodologies that use genetic similarity to identify groups of individuals with high (genetic) affinity or likeness [13]. The aim is to identify relatively homogenous groups of individuals that approach populations consistent with a Hardy-Weinberg model and are resultantly more appropriate for many of the assumptions built into many of the methods used in genomic epidemiology studies [14,15]. We leverage public data from the 1000 Genomes Project (1KG) [16] to provide reference populations from four, therein described, superpopulations or (sub)-continental ancestry groups (CAGs) – namely, Africa (AFR), Europe (EUR), South Asia (SAS), and East Asia (EAS). We note that we will refer to the groupings or clusters of individuals derived by this work, not as populations, but as groups or clusters of individuals. Further, the groups and clusters identified here are used as discrete units, but ancestry does not have decisive boundaries and is a continuum [17–20]. The use of discrete units is an analytical simplification. Finally, the overarching purpose of our study is to provide a description of the population structure present in the UKBB as an aid to future research investigating the health of individuals from diverse ancestries.

## Results

### Estimations of continental ancestry

Each of the 78,296 UKBB “non-white British” were included in a supervised ADMITXTURE analysis to estimate a proportion of ancestry to each of African (AFR), European (EUR), South Asian (SAS), and East Asian (EAS) continental ancestry groups (**Figure 1**). The proportion of continental ancestry is further illustrated, for each individual, within the context of UKBB population structure on principal components (PC) one and two as provided by the UKBB (**Figure 2**). AFR ancestry (**Figure 2A**) runs largely parallel with PC1, the major axis of variation. EUR ancestry runs at a roughly 135-degree angle (**Figure 2B**) along PC1 and PC2, while SAS (**Figure 2C**) and EAS (**Figure 2D**) ancestry run, largely, along PC2. Of the approximately 78,000 UKBB samples included in the ADMIXTURE analysis 50,685, 6,653, 2,782, and 2,364 individuals had 80% or more of their ancestry attributed to the EUR, AFR, SAS, and EAS continental super-populations, respectively. These individuals were carried forward into further analyses of population structure within these continental ancestry groups (CAGs). The 80% threshold was chosen to allow some error in the broader continental classification whilst also placing a limit on the complex structure and admixture evaluated in these subsets. A total of 15,812 “non-white British” UKBB study participants were not included in any of the four CAGs, given the methods and cut-offs used here.

**Figure 1.**
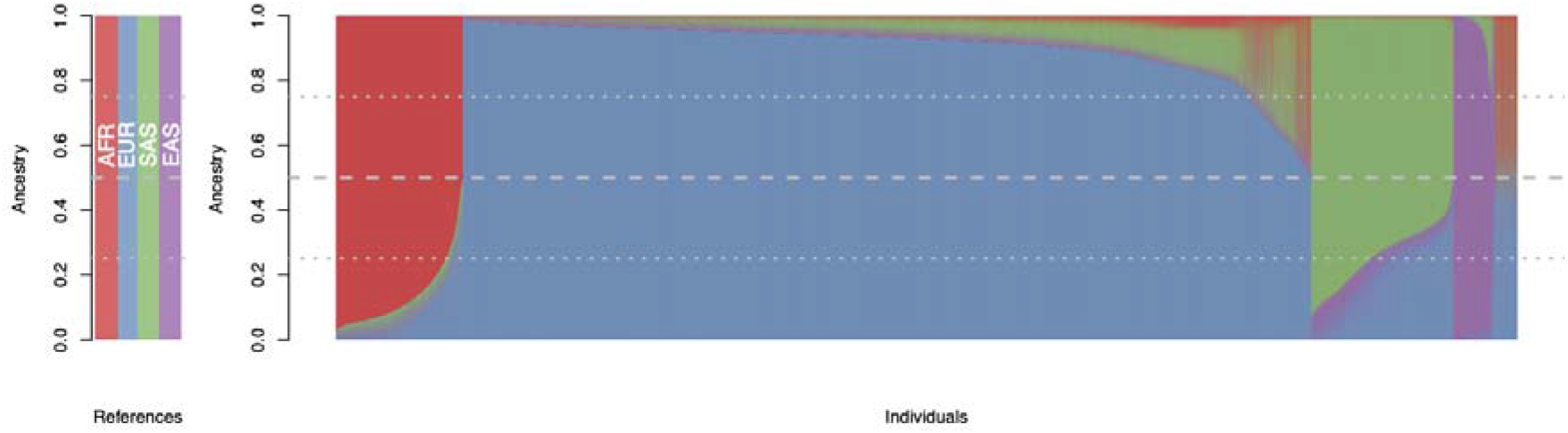
Ancestry estimates for the UKBB non-white British subset: Estimates of ancestry proportions for each UKBB participant previously labeled as non-white British individuals by UKBB. Ancestry was derived from a supervised ADMIXTURE analysis using four 1000 Genomes reference populations - Yoruba in Ibadan, Nigeria for (AFR) Africa, British in England, and Scotland for (EUR) Europe, Indian Telugu in the UK for (SAS) South Asia, and Han Chinese South for (EAS) East Asia.

**Figure 2.**
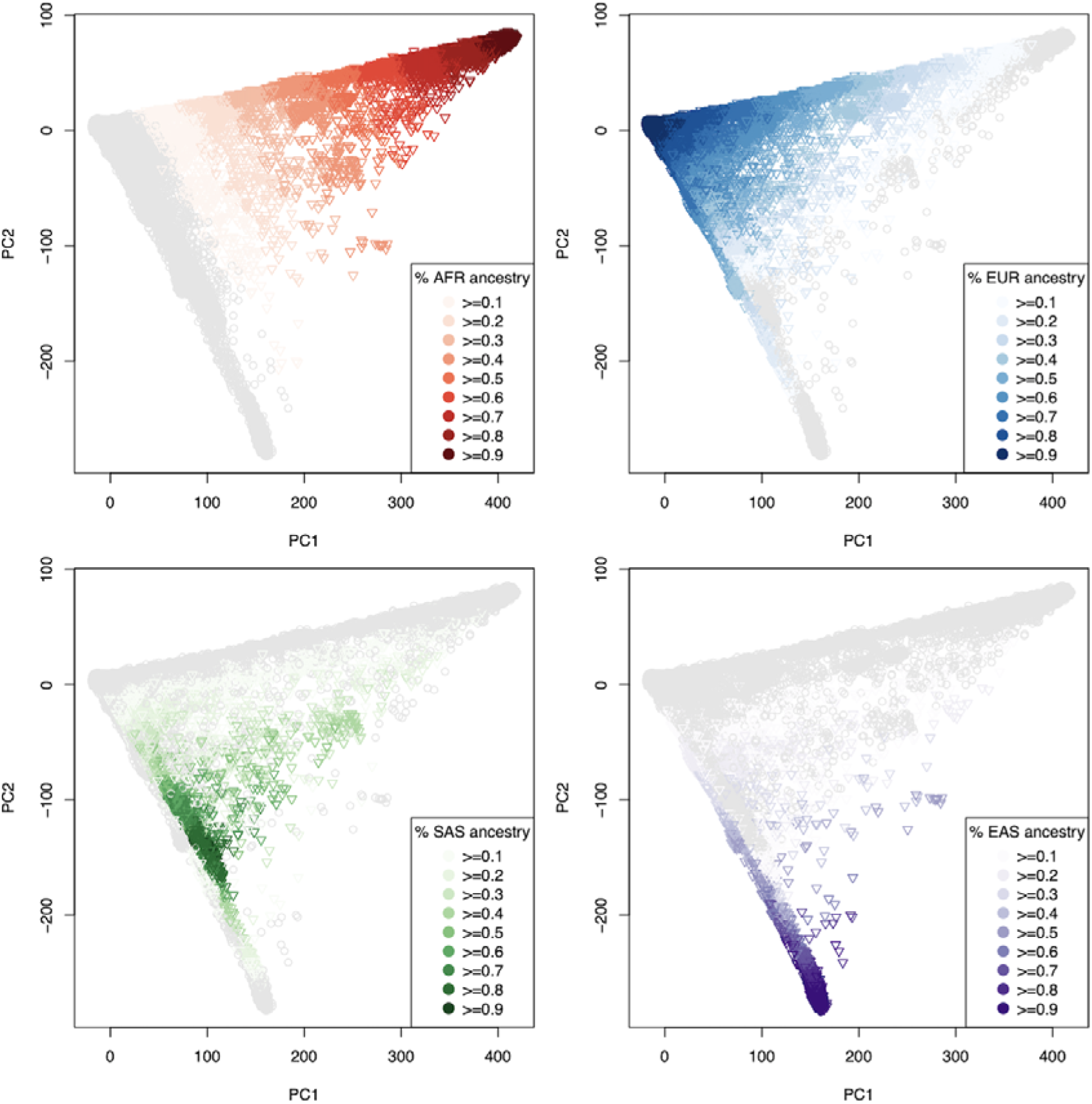
Ancestry proportions on UKBB PCs: Continental (A) African, (B) European, (C) South Asian, and (D) East Asian ancestry proportions placed on principal components one and two, as supplied by the UK Biobank.

### Population structure within continental regions

To evaluate the level of population structure among the UKBB CAGs, we first re-estimated principal components for each, while also projecting individuals from 1KG populations from each super-population respectively, onto the newly derived PCs (**Figure 3**, **Supplementary Table 1**). For each there is considerable overlap between UKBB individuals and 1KG populations, providing some context for the diversity that is present within the UKBB. In the AFR continental ancestry group principal component one distinguishes West African from East African 1KG populations, while PC3 distinguishes among populations of West Africa (**Figure 3A**). In the EUR continental ancestry group, the PCs and 1KG populations illustrate a strong North-South axis along PC2, with a similar but less distinctive trend on PC1 (**Figure 3B**). In the SAS continental ancestry group, there is a South-North trend along PC1, but no remarkable pattern can be attributed to the PCs (**Figure 3C**). The 1KG sample populations in the EAS ancestry group appear to indicate a North-South axis along PC1, and a West to East axis along PC2 (**Figure 3D**).

**Figure 3.**
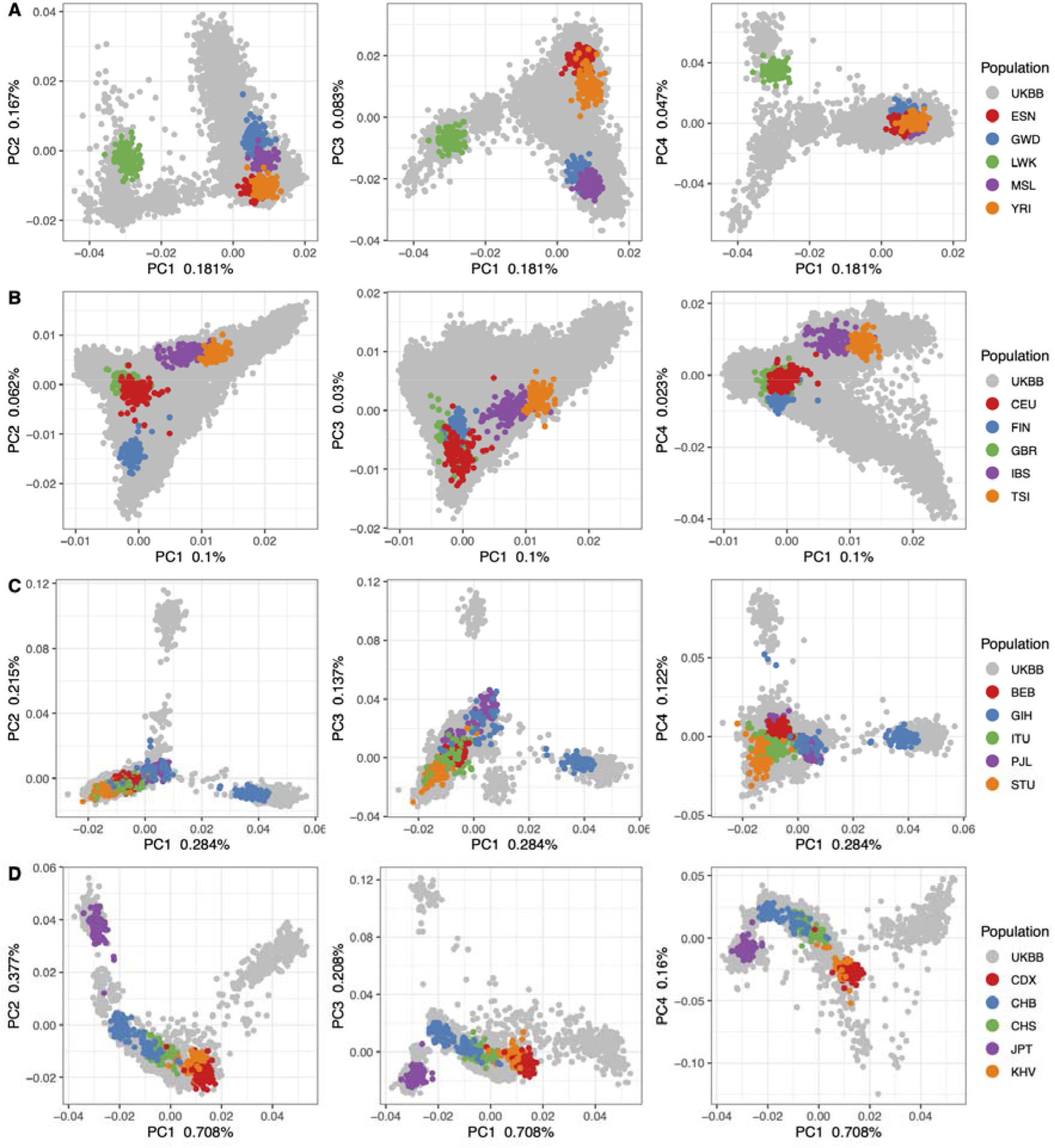
UKBB continental PCs with 1000 Genomes populations: Principal components one through four for each CAG. UKBB samples are colored in gray, while the 1KG sub-populations for each CAG are plotted in other colors, as indicated by each legend. The proportion of variation explained by each PC is indicated on each axis.

### K-means clustering of PCs

Given that many population genetics and epidemiological analyses, such as genome-wide association studies, depend on limited population structure, a common desire is to have a relatively homogeneous population sample for these analyses. As such, we used an unsupervised algorithm to identify groups of individuals that approach Hardy-Weinberg population assumptions. To do so we performed a K-means analysis on the top PCs (see Methods, **Supplementary Figure 1**), from each CAG, to identify ‘K’ subclusters or groups within each. An optimum number of K-clusters was determined by a silhouette analysis (see Methods, **Supplementary Figure 2**). For each CAG, using only the UKBB participants, we identified seven, two, four, and three K-clusters of individuals for AFR, EUR, SAS, and EAS, respectively (**Supplementary Figure 3**). However, for the EUR CAG we choose the second-best K-cluster (K=6) for the remaining analyses to improve our ability to investigate the utility of this analytical method to discriminate population structure (**Figure 4**).

**Figure 4.**
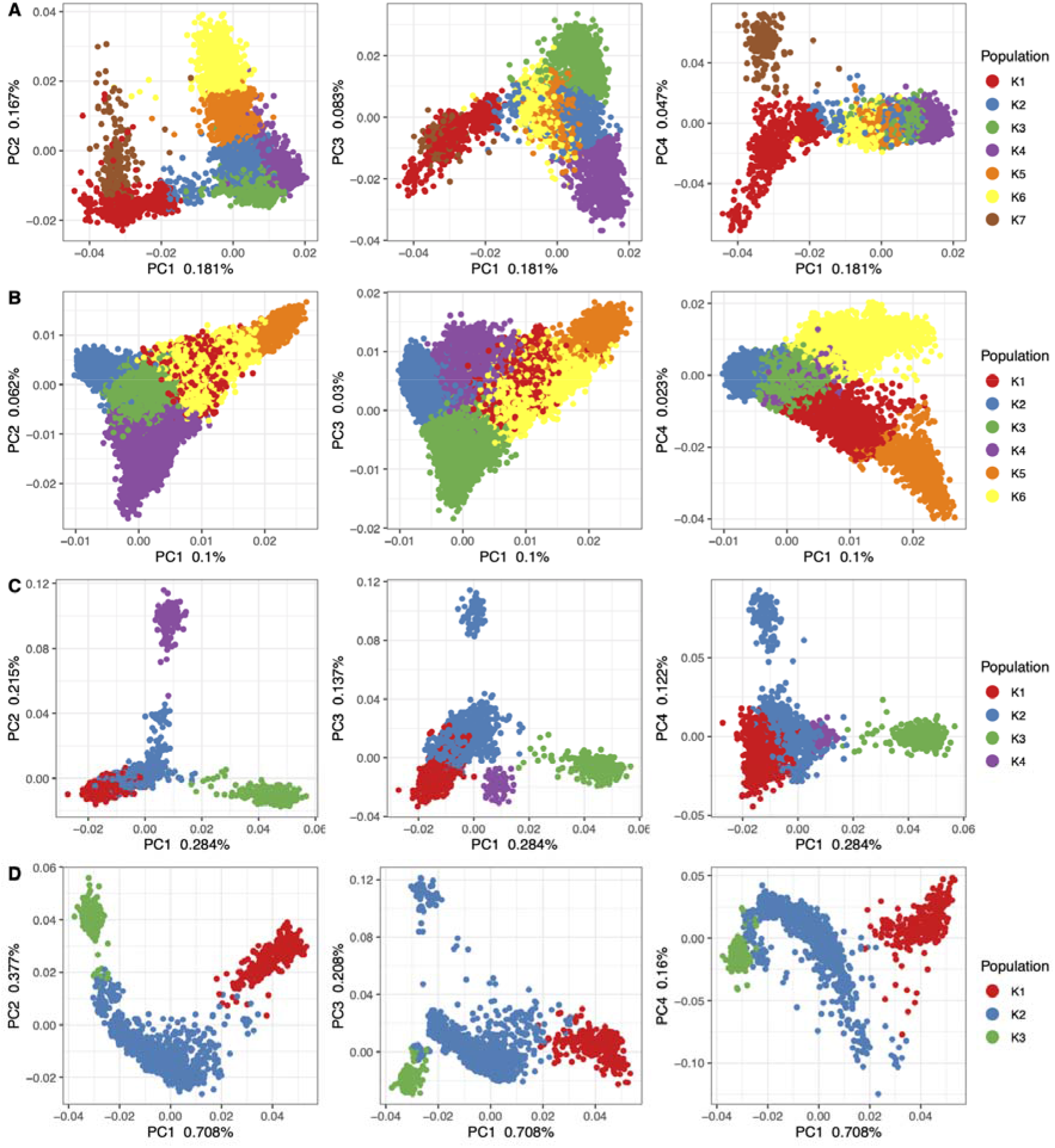
UKBB continental PCs with K-means clusters: Principal components one through four for each CAG with each individual colored by its assigned K-means population cluster, as indicated by each legend. The proportion of variation explained by each PC is indicated on each axis.

### Country of birth

To evaluate the informativeness of these K-clusters we mapped each individuals’ country of birth and United Nations (UN) geographic regions onto the PCs (**Figure 5** and **Supplementary Figure 4-5**). These figures further illustrate the diversity and structure present in the sample. Each CAG presents an observable degree of population structure, and region of birth (ROB) data illustrates non-specific associations between CAGs and ROB (Figure 5). For example, a large number of individuals have an East African ROB but are estimated to have more than 80% of their ancestry from South Asia (Figure 5 C and G). Nevertheless, ROB data illustrates structure across principal components for each CAG. Yet to ascertain if there is a correlation among the K-clusters identified above and the self-reported place of birth we performed a correspondence analysis for each CAG. The analyses indicate a correlation between K-means clusters and the UN regions for each continent: AFR (Dim1 53.29%, Dim2 41.88%), EUR (Dim1 58.25%, Dim2 28.67%), SAS (Dim1 80.00%, Dim2 18.2%), EAS (Dim1 92.11%, Dim2 7.89%) (**Figure 6A**). When UN regions for a smaller geographical region were substituted, namely country of birth (COB; Supplementary Figures 6-9), an attenuated but correlated structure remained: AFR (Dim1 28.32%, Dim2 25.02%), EUR (Dim1 40.43%, Dim2 31.89%), SAS (Dim1 61.60%, Dim2 25.31%), EAS (Dim1 50.49%, Dim2 49.51%) (**Figure 6B**, **Supplementary Figure 10**).

**Figure 5.**
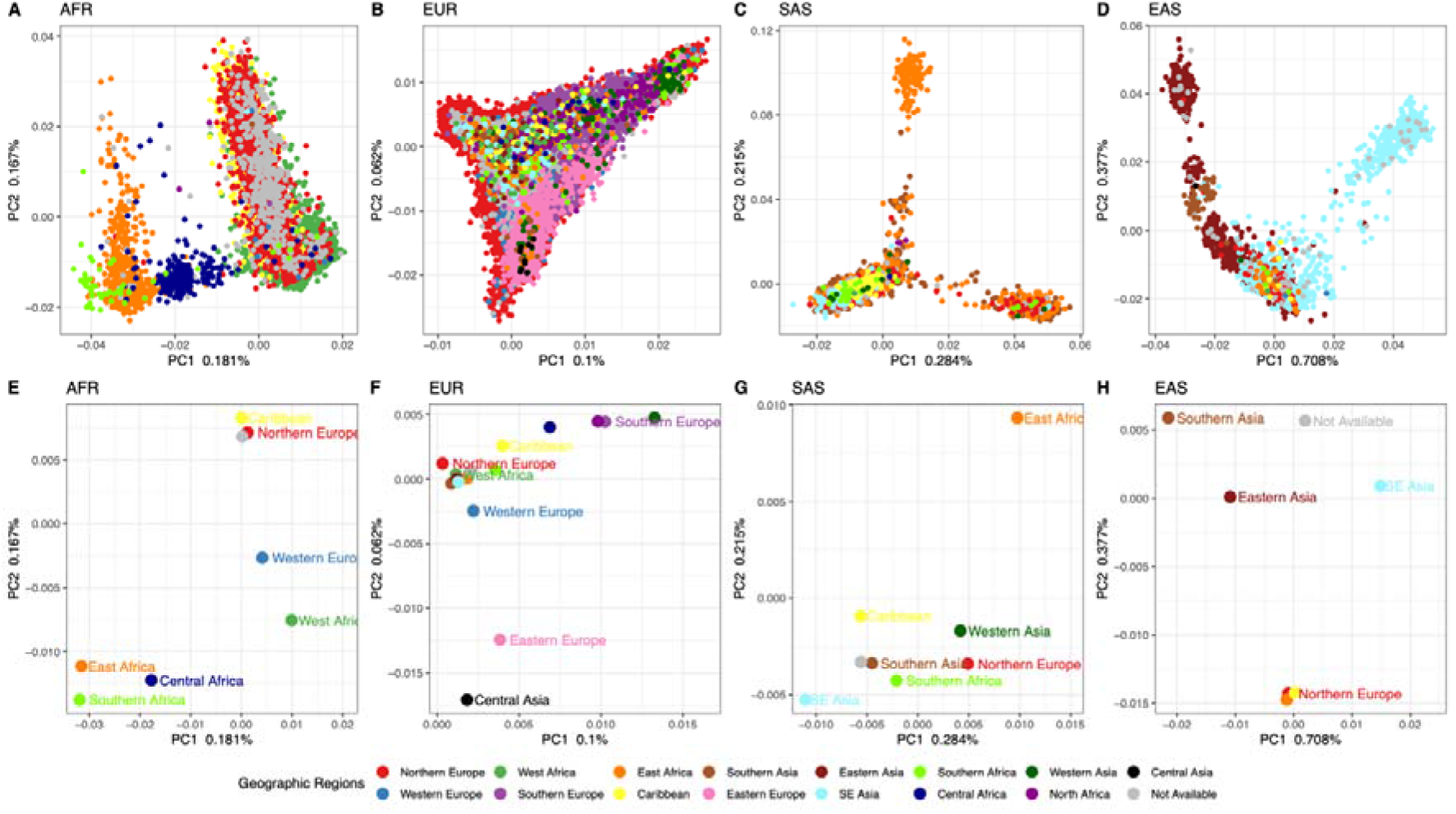
Principal components for CAG with geographic regions of birth: Principal components one and two for each CAG, with (A-D) individuals colored by their region of birth (A-D), and with (E-H) the PC center also colored by region of birth. PC centers were estimated as the average PC1 and PC2 values for all individuals of that ROB. Regions of birth are denoted in the figure legend, and the proportion of variation explained by each PC is indicated on each axis.

**Figure 6.**
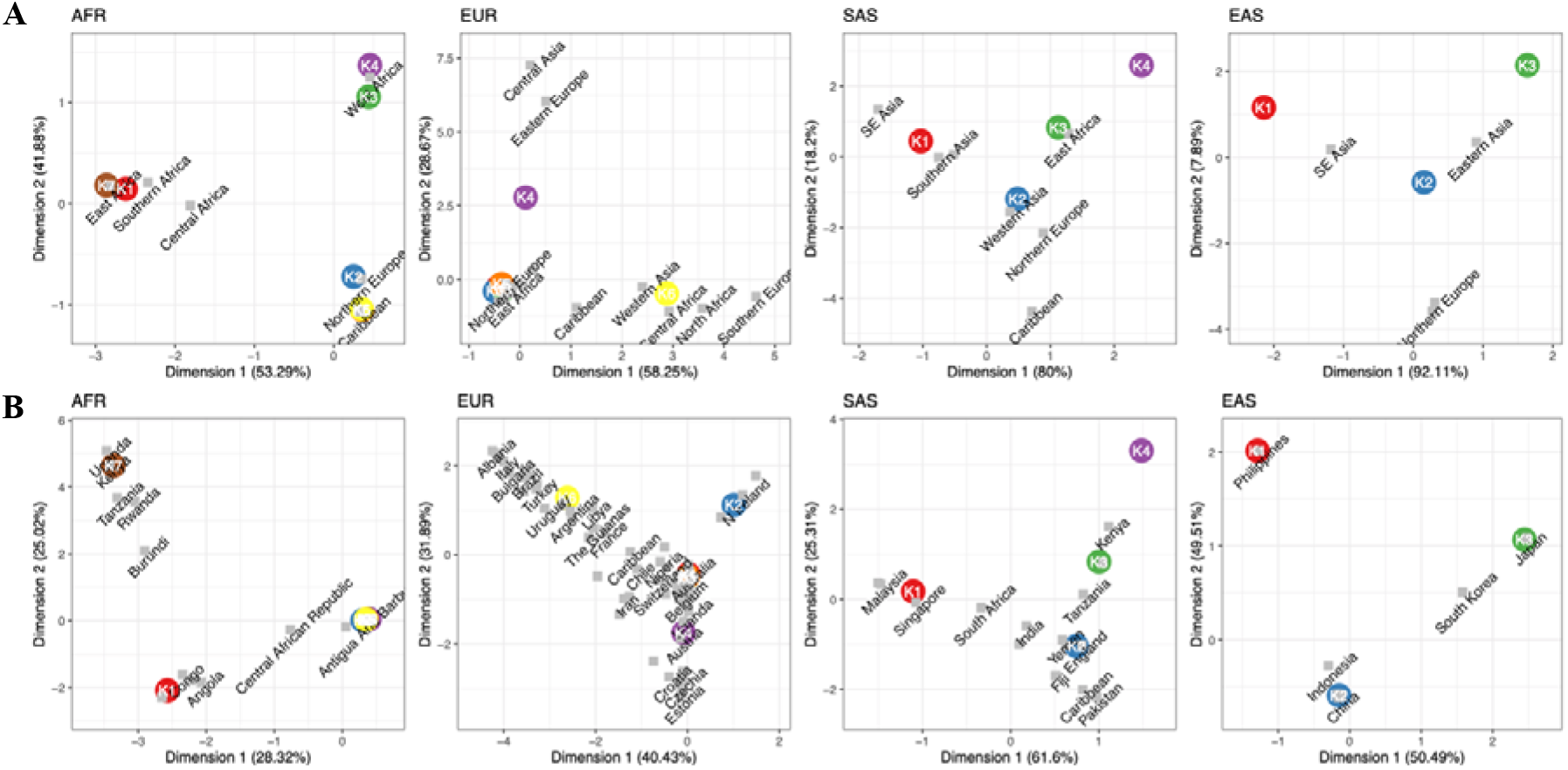
Correspondence analysis: Correspondence plots between (A) K-means population clusters (colored circles) and regions of birth (grey squares), and (B) K-means population clusters (colored circles) and country of birth (grey squares) (B). The x and y axes are the first and second dimension of each correspondence analysis, respectively, with the proportion of variance explained indicated in the parentheses of each axis.

### Population differentiation

An evaluation of the degree of population differentiation within each CAG was performed by estimating Fst, or the fixation index between each pair of K-cluster groups and 1KG populations. All single-nucleotide polymorphisms (SNPs) that were included in each CAG’s principal component analysis were used here. An average, minimum, and maximum estimate was used to summarize the distribution of estimates between pairs (**Figure 7**). Relative to the population differentiation observed in the 1KG sample populations we observed, on average, a small degree a population differentiation among AFR and EUR K-means clusters, and larger average estimates among SAS and EAS groups. Among the UKBB samples average Fst estimates indicate that the EAS CAG has the largest amount of population differentiation with an average Fst of 0.0133. This is followed by SAS with an average estimate of 0.0092, EUR with 0.0037, and finally AFR with the smallest average estimate of 0.003. However, we note that these estimates were derived from SNPs with a European ascertainment bias and as such they may not coincide with analyses using an unbiased set of genetic variants.

**Figure 7.**
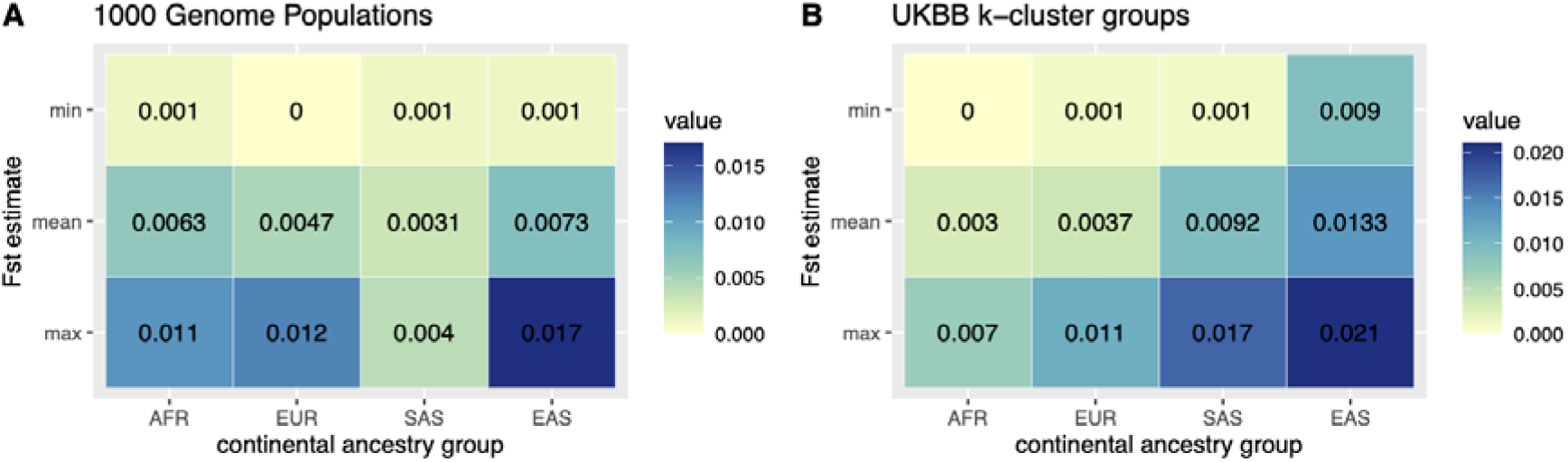
Fst estimates: The minimum, mean and maximum fixation index values for each CAG in the 1KG project and the UK Biobank dataset. Fst values in the 1KG project are between the sub-populations of each super-population, while UK Biobank estimates are derived between K-means population cluster of each CAG.

## Discussion

Here we present an analytical pipeline to identify individual participants of the UKBB study with diverse and under-represented ancestries to be used in genomic epidemiology studies. Whilst cohort studies centred in diverse geographic locations are essential for elucidating the effect of environment and genotype on disease, the diversity present in deeply phenotyped studies such as the UKBB should be utilized where possible. This study presents a description of some of the diversity present in the UKBB. Further, the methods presented here provide an approach to identify subsets of individuals to help broaden, inform, and improve the relevance of genetic epidemiological studies and their findings for those of, in this specific instance, a non-white British ancestry (**Figure 8**).

**Figure 8.**
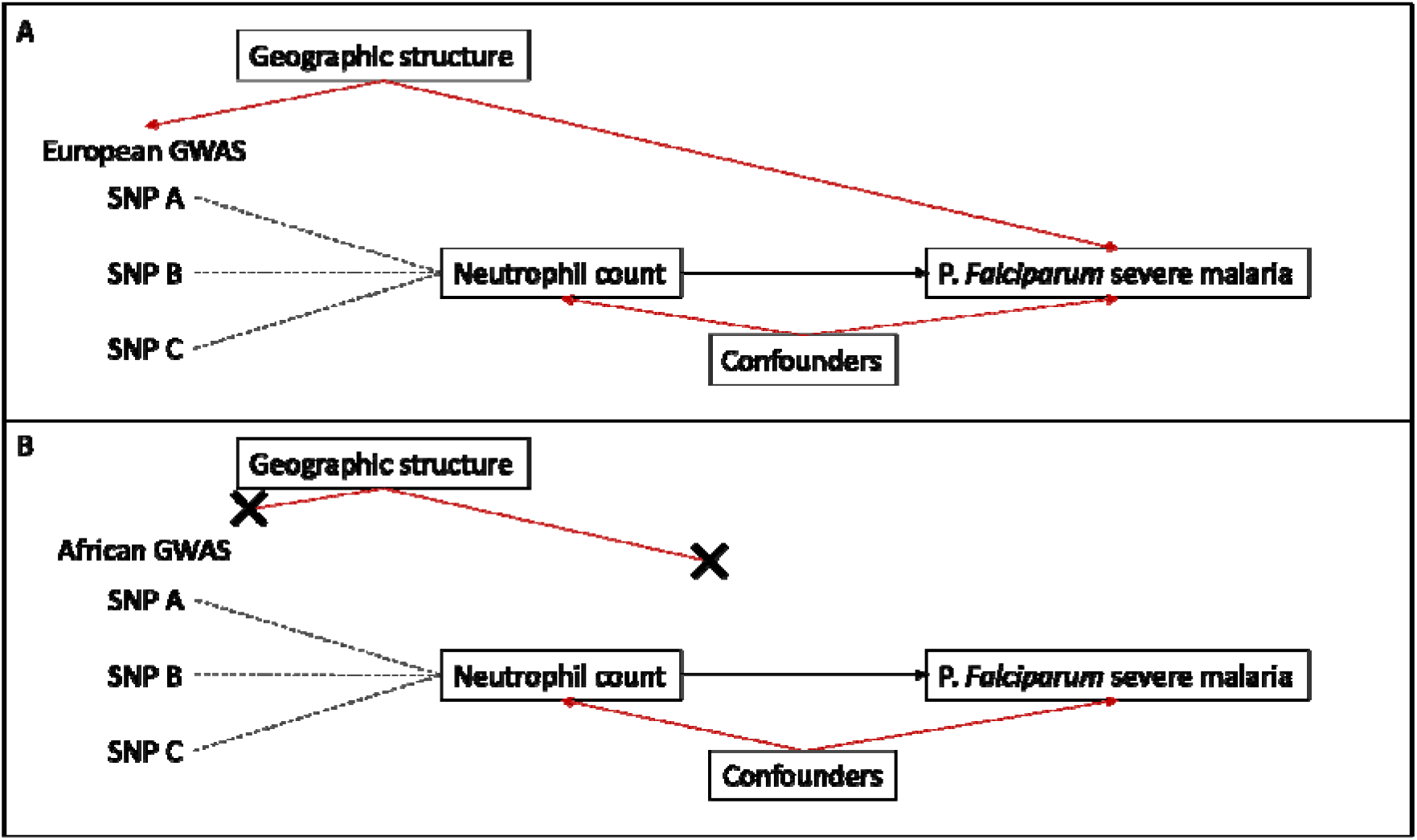
Graph outlining the possible effects of geographic structure in population genetics: Suppose one might want to use Mendelian randomization to study the relationship between neutrophil count and severe malaria caused by P. *Falciparum* – a disease largely absent in European environments. Using summary statistics from a neutrophil count GWAS derived from individuals with European ancestry (Box 1A) may affect estimates due to geographic structure (Ancestry + Demography + Environment). This can be overcome by running a GWAS in people of African ancestry (Box 1B).

Throughout the paper, when we speak of ancestry, we are referring to “genetic ancestry”, or individuals who share a demographic history [13,21,22]. They would, at the population level, share a history of mutation, genetic drift, recombination, migration, natural selection, environment, and culture (niche construction). As a product, they would have different genetic variants, allele frequencies, and patterns of linkage disequilibrium across their genomes [23–25].

The need to perform analyses, like association studies, separately in unique ancestral populations is largely born from the need to avoid correlations between phenotype and genetic ancestry, or differences in allele frequencies among populations [13,26]. For example, if a disease (or environmentally influenced trait) is more frequent in ancestral population ‘A’ than it is in ‘B’ and your association analysis pools these ancestral populations together you may erroneously identify any allele that is more frequent in population ‘A’ as a genetic variant associated with the disease. To avoid these confounding issues, analyses are commonly limited to relatively homogenous populations.

In genome-wide association studies, the aim is to derive accurate unbiased effect estimates for a genetic variant on a trait. However, the task becomes increasingly challenging, as variation in genetic ancestry comes with different allele frequencies, genetic backgrounds and environments [27]. Methods such as the inclusion of relatedness matrixes and principal components [28–31] are used to account for cryptic relatedness and undetected, fine-scale population stratification. In addition, they are also used to account for correlations between phenotype and genetic ancestry [32,33]. However, are the inclusion of relatedness matrixes or principal components enough to control the structure present in the CAGs presented here? Or would smaller (K-means clusters) more homogenous populations be better suited to epidemiological analyses, like GWAS?

The problems introduced by population stratification persist even in populations like the “white British” subset of the UKBB, where individual genetic variants and polygenic scores for individual traits can retain correlations with geography, even after correcting for population structure [34,35]. Moreover, when sampling populations across Europe – where genetic ancestry does mirror geography [36,37] – and meta-analysing independently run GWASs [38], effect estimates appear to retain a bias introduced by population structure [39,40]. These fine scale issues exemplify some of the reasons for performing separate epidemiological analysis, like GWAS, for populations with deeper population differentiations, i.e. unique ancestries, demographic histories, and environments.

The complications of population stratification and opportunities for improving health outcomes for more people, even at the continental level, are precisely why a description of the structure within each continental ancestry group was provided here. Namely, the structure present within a CAG, as identified here, may also be too great to be properly accounted for with common methodologies and may thus need to be resolved into smaller more homogenous groups. At the very least, careful consideration is warranted when interpreting results where CAGs are used - because structure matters [41]. The unsupervised clustering performed within each CAG is not a perfect solution for identifying true “populations” – an exercise that may in fact be an impractical goal – but it is a method to identify groups of individuals with a more similar, homogeneous ancestry. Other techniques like uniform manifold approximation and projection [42] or more explicit leveraging of self-described ethnicity could help improve the identification of homogenous groups. Self-described ethnicity is not a synonym for genetic ancestry though, as it is a sociocultural construct. It would however help inform cultural, social, and other environmental influences – important aspects of a “population” - on phenotypes and disease [22].

In summary, we assigned individuals to continental ancestry groups (**Figure 1 and 2**); illustrated the structure present among individuals within each CAG (**Figure 3**), identified unsupervised clusters or groups of individuals within each (**Figure 4**) and demonstrated that those clusters have an affinity to regions and countries of birth – i.e. the K-means clusters are consistent with geographic structure and isolation by distance models [43,44] (**Figure 5**). Notably, each CAG presents extensive structure, inconsistent with a randomly mating population, but rather with the sampling of unique, geographically distant populations. In particular, East Asian, South Asian, and African CAGs have isolated, or discontinuous groups of individuals in the UKBB sample, exemplified in the K-means clustering analysis (Figure 4) [19,20]. For example, groups K1 and K3 in the EAS CAG (**Figure 4D**) epitomizes this discontinuous structure as they correspond to individuals born on the islands of Philippines and Japan, respectively (**Figure 5**, **Supplementary Figure 8**).

The methods employed here do have several limitations: First, a single 1KG population was used to represent each of four continental ancestry groups evaluated – Africa, Europe, South Asia, and East Asia. One population is a poor proxy for all of the variation present in any one (sub)-continent. However, as the 1KG project does not have optimal population coverage, including more or all the 1KG populations of a CAG would still poorly represent all the variation present in a (sub)-continent and would complicate the assignment of individuals to a single ancestry group. Second, our analysis was limited to four (sub-)continental ancestry groups, to the exclusion of the Americas (AMR, a 1KG superpopulation). Populations from the Americas often have a large and varying amount of recent admixture from various European and African populations [25,45–49]. As such, including an AMR population in the ADMIXTURE analysis, as a reference population, could confound the genetic ancestries being estimated. However, whilst we limit this study to a few, broad, well characterized ancestry groups the approach presented her can be generalised to other, specific ancestries.

Third, the UKBB Axiom array used to genotype all UKBB participants was designed to optimize imputation of a European population while also including genetic variants previously associated with disease and other phenotypic traits derived from studies primarily conducted in European populations [11,12]. As a product, the genomic data used here will have an ascertainment bias [50] that would influence imputation accuracy (although no imputation data was used here), allele frequency distributions, estimates of linkage disequilibrium and diversity and divergence within and among populations. Each of these may influence estimations of population differentiation, principal component estimates and the inferences made from them [51,52]. Specific study designs [53,54] have been made to remove ascertainment bias in genotype arrays so that unbiased inferences could be made for a wider range of genetic ancestries, but this was not available here.

Fourth, the principal components illustrated and used in the unsupervised K-means clustering analyses were derived from the UKBB participants only and resultantly represents the diversity (point three) and genetic ancestry found in that data set. The inclusion or use of other public data sets with more numerous sample populations, that better represent regional, or continental diversity will provide alternative patterns of structure. Fifth, we are limited by the reference population used in the analyses. Whilst the 1KG data set shall remain an essential reference panel for broad analyses like those conducted here, researchers with specific continental or geographically specific research questions could strengthen and refine the observations made here by including other geographically specific data sets. Finally, the unsupervised K-means clustering analysis is dependent upon the number of PCs included in it. Here the number of PCs chosen did have an element of subjectivity (Supplementary Figure 1). Whilst analytical methods are available to select a number of informative PCs [55], we did not implement such methods here. Given that the K-means algorithm weights each PC equally, we sought to limit the PCs included to only those with the largest proportions of variance explained and not necessarily all that are analytically estimated to be informative.

## Conclusions

The approach presented here demonstrates a method to leverage the deeply phenotyped and widely used UKBB data set to help improve the inclusion and equity of epidemiological studies for under-represented populations. Careful considerations must be given to the diversity present within continental ancestry groups. However, given the thousands of individuals present in the genetic ancestry groups identified here, the UKBB data set shall prove insightful for studies of health and disease in populations beyond the British Isles. While the methods presented here do not describe a perfect solution to identify populations, we hope that they provide an avenue to leverage the diverse data available in UKBB and a methodological platform to improve and build upon.

## Methods

### Description of working environment

All analyses were performed in a Linux environment supported by the University of Bristol’s Advanced Computing Research Centre (ACRC) using the following publicly available software packages: Plink v1.9 and v2.0 [56,57], ADMIXTURE v1.3.0 [58,59], and EIGENSOFT v8.0.0 [29,30]. In addition, bespoke scripts, analyses, and figures were run and generated in the R environment using version 3.6.2 on the ACRC computer clusters and version 4.0.2 (Taking Off Again) on local computers [60].

### UK Biobank data

This research has been conducted using the UKBB Resource under Application Number 15825, from which directly genotyped SNP data (N=784,256 SNPs) were made available. It includes data for a total of 78,296 individuals identified by UKBB as “non-white British” participants – our analyses were restricted to this subset. In addition to genotypic data, we also acquired several variables of interest (self-described ancestry, country of birth) data for this subset of individuals. 365 exclusions were made when filtering those with sex chromosome mismatch and/or aneuploidy, and outliers with high genetic heterozygosity and missing rates [61].

### 1000 Genomes data

Genetic data (v5a.20130502) from phase three of the 1KG, which includes data from 5 continental, or 1KG described super-populations [Europe (EUR), East Asia (EAS), South Asia (SAS), Africa (AFR), and the Americas (AMR)], were used to provide reference populations for admixture analyses and population structure inferences ([62] http://ftp.1000genomes.ebi.ac.uk/vol1/ftp/). Our analyses did not include populations from the AMR superpopulation. This is to maintain a simplified analysis that avoided the complicating factors of the potentially recent admixture events that occurred in the Americas. Included in our analyses are five populations from 1KG super-population label: (AFR), also known as the continental Africa ancestry group (1) Yoruba in Ibadan, Nigeria (YRI); (2) Luhya in Webuye, Kenya (LWK); (3) Gambian in Western Division, The Gambia - Mandinka (GWD); (4) Mende in Sierra Leone (MSL) and (5) Esan in Nigeria (ESN). Five populations from the super-population label EUR or the continental Europe ancestry group: (1) Utah residents with Northern and Western European ancestry (CEU); (2) Toscani in Italia (TSI); (3) British in England and Scotland (GBR); (4) Finnish in Finland (FIN) and (5) Iberian populations in Spain (IBS). Five populations from the super-population label SAS or the continental South Asian ancestry group: (1) Gujarati Indian in Houston, Texas (GIH); (2) Punjabi in Lahore, Pakistan (PJL); (3) Bengali in Bangladesh (BEB); (4) Sri Lankan Tamil in the UK (STU) and (5) Indian Telugu in the UK (ITU). Finally, five populations from the super-population label EAS or the continental East Asian ancestry group: (1) Han Chinese in Beijing, China (CHB); (2) Japanese in Tokyo, Japan (JPT); (3) Han Chinese South (CHS); (4) Chinese Dai in Xishuangbanna, China (CDX) and (5) Kinh in Ho Chi Minh City, Vietnam (KHV).

### Merging UK Biobank and 1000 Genomes

The directly genotyped data from UKBB was used to identify SNPs with the same SNP identifier (RefSNP ID) present in the 1KG data set. A total of 718,711 SNPs were identified with the same ID and extracted from both data sets using PLINK v2.0. The two datasets were then merged using the -bmerge function in PLINK v2.0. After removing problematic SNPs (e.g. multi-allelic, duplicate) in the merge step, a total of 718,487 SNPs remained.

### Linkage disequilibrium pruning

Prior to ancestry estimation the merged dataset was reduced to a set of independent SNPs based on linkage disequilibrium (LD) estimates using the PLINK v2.0 function and parameters “--indep-pairwise 50 10 0.025”, indicating an r^2^ threshold of 0.025, a window size of 50 kilobases and a window step size of 10 kilobases. In addition, 24 previously identified genomic regions with extensive linkage disequilibrium were also excluded [63,64]. LD estimates in this analysis were limited to unrelated individuals from the 1KG YRI population sample. A total of 30,320 SNPs remained following LD pruning.

### Estimating African, European, South Asian, and East Asian ancestry

Four 1KG populations were included as reference populations in a supervised Admixture (v1.3.0) analysis. They were (1) British in England and Scotland (GBR), of the European ancestry (EUR) superpopulation, (2) Yoruba in Ibadan, Nigeria (YRI), of the African ancestry (AFR) superpopulation, (3) Indian Telugu in the UK (ITU), of the South Asian ancestry (SAS) superpopulation, and (4) Han Chinese South (CHS), of the East Asian ancestry (EAS) superpopulation. These singular population samples were chosen to broadly represent each of their four respective continental (superpopulation) ancestry groups, with an average population differentiation (Fst, or fixation index) value of 0.1055 amongst them, as estimated by ADMIXTURE. The supervised ADMIXTURE analysis provides, for each UKBB sample, a proportion of ancestry for each of the four reference populations. Those individuals with at least 80% of their ancestry attributed to one continental ancestry group, or 1KG defined superpopulation, were carried forward into further analyses.

### Derivation of continental principal components

Unrelated individuals in each CAG, including both 1KG and UKBB samples with >=80% ancestry to that CAG were identified (using all 718,487 SNPs in the overlapping data set, and the plink (v1.9) function --rel-cutoff and a minor allele frequency (MAF) filter of 0.05 (--maf 0.05)). Then for each CAG and using all (1KG + UKBB) unrelated individuals assigned to the CAG, a list of approximately 40 thousand LD independent SNPs were identified (using the plink (v2.0) function --indep-pairwise 50 10 0.025 (--indep-pairwise 50 10 0.02 for AFR and --indep-pairwise 50 10 0.05 for SAS) along with a MAF filter of 0.01, and the exclusion of the 24 previously identified genomic regions with extensive linkage disequilibrium [63,64]). New plink files including only the LD independent SNPs identified in step two were subsequently generated. smartrel from the EIGENSOFT (https://github.com/DReichLab/EIG) package was used to generate a new list of related individual pairs, along with our script “greedy_unrelated_selection.R” to identify a list of related individuals to exclude from principal component derivation [29,30]. An exception this step was made for the European CAG as its sample-size was prohibitively large to run smartrel, instead the list of unrelated individuals generated from step one was used. Finally, smartpca of the EIGENSOFT package was used to estimate principal components (PC), using only unrelated UKBB samples. Related and 1KG samples were subsequently projected upon these PCs by smartpca. Sample outliers were excluded from the PC analysis by smartpca with the following parameters: using 10 PCs to identify outliers (numoutlierevec), at six standard deviations from the mean (outliersigmathresh), and with 5 outlier removal iterations (numoutlieriter). **Supplementary Table 1** provides numbers for each of these steps, for each CAG. The EUR CAG was treated uniquely due to its larger sample-size. Smartpca was run twice as described above, once with “fastmode=NO” and then with “fastmode=YES”. The former provided estimates of the eigenvalues but not the eigenvectors, while the latter provided eigenvectors but not eigenvalues.

### K-means clustering of principal components

For each CAG, we estimated the variance explained by each principal component (PC) by dividing the eigenvalue of each PC by the sum of all eigenvalues. To identify the number of top PCs we generated a scree plot, using the variance explained estimates, and identified the elbow or valley in each plot (**Supplementary Figure 1**, **Supplementary Table 2**). The top PCs, and the top PCs only, were then used in an unsupervised K-means clustering analysis (k set from 2 to 20; using the function “kmeans()” from the R stats package) to identify clusters of UKBB individuals that maximize between cluster sums of squares and minimize within cluster sums of squares. An optimum number of clusters (k) was identified by silhouette analysis using the function “pamk()” from the fpc R package (**Supplementary Figure 2**) [65]. These analyses are implemented in our function “DetermineK()” found in this study’s GitHub repository.

### Correspondence analysis

Each UKBB study participants’ country of birth information was placed into United Nations defined geographic regions (**Supplementary Table 3**). To determine if the K-means population clusters have any relationship with an individual’s country of birth or country of birth UN-region we performed correspondence analyses (CAs) using the function “ca()” from the R package “ca”, for each continental ancestry group [36]. In addition, a chi-square test was performed on the contingency table used in the correspondence analysis. Any UN-region or country of birth with fewer than 10 observations was excluded. Individuals for which country of birth information was not available were also excluded.

### Population differentiation among K-means population clusters

For each CAG, we took the best K-means population clusters, as defined by the silhouette analysis, and re-ran smartpca. However, on this run we had smartpca provide for us only an estimation of the average fixation index (Fst) for each pair of populations in the data set, including 1KG populations and UKBB K-means clusters. This was done with the inclusion of the paramaters “fstonly” and “phylipoutname” [55], the latter of which provides a distance matrix of mean Fst values between populations. Estimations of Fst, which range from 0 to 1, provide a measure of population differentiation among populations. In brief, these describe the proportion of total variation at a SNP that is explained by variation between populations. For any SNP a value of 0 would indicate that minimal variation is attributable to variation between populations. A value of 1 would indicate a fixed difference i.e., the two populations are both invariable but for alternative alleles.

## Declarations

### Ethics approval and consent to participate

UK Biobank received ethical approval from the NHS National Research Ethics Service North West (11/NW/0382; 16/NW/0274) and was conducted in accordance with the Declaration of Helsinki. All participants provided written informed consent before enrolment in the study.

### Consent for publication

All authors consented to the publication of this work.

### Availability of data and material

Genetic data from UK Biobank were made available as part of project code 15825. Analytical code is available on GitHub at https://github.com/andrewcon/popgen-biobank.

### Competing interests

None to declare.

### Funding

AC acknowledges funding from a Medical Research Council PhD studentship (MR/N013794/1). NJT and REM acknowledge funding from the Medical Research Council (MC_UU_00011/1). NJT is the PI of the Avon Longitudinal Study of Parents and Children (Medical Research Council & Wellcome Trust 217065/Z/19/Z) and is supported by the University of Bristol NIHR Biomedical Research Centre (BRC-1215-2001). EEV, CJB, NJT and DH acknowledge funding from the Wellcome Trust (202802/Z/16/Z). EEV, CJB and NJT also acknowledge funding by the CRUK Integrative Cancer Epidemiology Programme (C18281/A29019). EEV and CJB are supported by Diabetes UK (17/0005587) and the World Cancer Research Fund (WCRF UK), as part of the World Cancer Research Fund International grant program (IIG_2019_2009). JZ is supported by the Academy of Medical Sciences (AMS) Springboard Award, the Wellcome Trust, the Government Department of Business, Energy and Industrial Strategy (BEIS), the British Heart Foundation and Diabetes UK (SBF006\1117). JZ is funded by the Vice-Chancellor Fellowship from the University of Bristol and is supported by Shanghai Thousand Talents Program. BA acknowledges funding from the Medical Research Council (MR/R02149x/1). The funders of the study had no role in the study design, data collection, data analysis, data interpretation or writing of the report.

### Authors’ contributions

AC, DH, and REM conceived the idea for the paper. AC and DH conducted the analysis. All authors contributed to the interpretation of the findings. AC and DH wrote the manuscript. All authors critically revised the paper for intellectual content and approved the final version of the manuscript.

## Acknowledgements

We are grateful to the UK Biobank study and its participants. This research has been conducted using the UK Biobank resource under Application 15825.

## Authors’ information (optional)

MRC Integrative Epidemiology Unit at the University of Bristol, Bristol, United Kingdom.

AC, REM, JZ, CJB, NJT, EEV and DH

Population Health Sciences, Bristol Medical School, University of Bristol, Bristol, United Kingdom.

AC, REM, JZ, CJB, NJT, EEV and DH

School of Translational Health Sciences, University of Bristol, Bristol, United Kingdom.

AC, CJB and EEV

School of Cellular and Molecular Medicine, University of Bristol, Bristol, United Kingdom.

BA

## List of abbreviations

1KG: 1000 Genomes Project
ACRC: Advanced Computing Research Centre
AFR: African
AMR: Americas
BEB: Bengali in Bangladesh
CA: correspondence analysis
CAG: continental ancestry group
CDX: Chinese Dai in Xishuangbanna, China
CEU: Utah residents with Northern and Western European ancestry
CHB: Han Chinese in Beijing, China
CHS: Han Chinese South
COB: country of birth
EAS: East Asian
ESN: Esan in Nigeria
EUR: European
FIN: Finnish in Finland
Fst: fixation index
GBR: British in England and Scotland
GIH: Gujarati Indian in Houston, Texas
GWAS: Genome-wide association study
GWD: Gambian in Western Division, The Gambia - Mandinka
IBS: Iberian populations in Spain
ITU: Indian Telugu in the UK
JPT: Japanese in Tokyo, Japan
KHV: Kinh in Ho Chi Minh City, Vietnam
LD: linkage disequilibrium
LWK: Luhya in Webuye, Kenya
MAF: minor allele frequency
MSL: Mende in Sierra Leone
PC: principal component
PJL: Punj abi in Lahore, Pakistan
ROB: region of birth
SAS: South Asian
SNP: single-nucleotide polymorphism
STU: Sri Lankan Tamil in the UK
TSI: Toscani in Italia
UKBB: UK Biobank
UN: United Nations
YRI: Yoruba in Ibadan, Nigeria

---

**Supplementary Figure 1 Continental ancestry PCA Scree plots:** Legend: Scree plots illustrating the proportion of variation explained by each of the top 20 PCs, in each UKBB continental ancestry principal component analyses. The Scree plots were used to identify the number of top PCs to carry forward into the K-means clustering analysis. The continental ancestry supergroups are Africa (AFR), Europe (EUR), South Asia (SAS), and East Asia (EAS). The number of PCs selected as top PCs are AFR = 4, SAS = 5, EAS = 4, EUR = 5. The horizontal line in each plot denotes where 10% variance explained is in each plot to aid in inter-CAG comparisons.

**Supplementary Figure 2 K-means k selection with silhouette analysis:** Selection of an optimum number of k clusters in the K-means analysis of the top PCs, by silhouette analysis. A silhouette plot for each UKBB continental supergroup (AFR) African, (EUR) European, (SAS) South Asian, and (EAS) East Asian is provided. The x-axis indicates the number of k clusters evaluated, and the y-axis provides an estimate of the average silhouette width (ASW). ASW is an estimation of cluster quality, or intra- and inter-cluster distances derived from a partitioning around medoids (PAM). The optimum number of k clusters in each UKBB continental supergroup were identified as AFR = 7, EUR = 2, SAS = 4, and EAS = 3.

**Supplementary Figure 3 UKBB continental ancestry group PCs with K-means clusters:** UK Biobank continental ancestry group PCs with K-means population clusters color coded: AFR (A), EUR (B), SAS (C), EAS (D).

**Supplementary Figure 4 Population structure by UN defined geographic region:** UK Biobank continental ancestry group PCs colored by the region of birth color coded: AFR (A-C), EUR (D-F), SAS (G-I), EAS (J-L).

**Supplementary Figure 5 Population structure centers, as defined by UN geographic region:** UK Biobank continental ancestry group PCs 1-4 with region of birth centers (averaged across all individuals from each ROB) colour coded: AFR (A-C), EUR (D-F), SAS (G-I), EAS (J-L).

**Supplementary Figure 6 Population structure by country of birth in AFR by region:** UK Biobank continental ancestry group PCs 1-4 for the AFR CAG divided by UN regions of birth: Eastern (A), Central (B), Western (C), Northern/Southern (D). Samples are color coded by their country of birth.

**Supplementary Figure 7 Population structure by country of birth in EUR by region:** UK Biobank continental ancestry group PCs 1-4 for the EUR CAG divided by UN regions of birth: Northern (A), Eastern (B), Southern (C), Western (D). Samples are color coded by their country of birth.

**Supplementary Figure 8 Population structure by country of birth in SAS and EAS:** UK Biobank continental ancestry group PCs 1-4 for the SAS and EAS CAGs divided by UN regions of birth: Southern Asia (A), East Asia and South-eastern Asia (B). Samples are color coded by their country of birth.

**Supplementary Figure 9 Population structure centers by country of birth:** UK Biobank continental ancestry group centers colored by the country of birth: AFR (A-C), EUR (D-F), SAS (G-I), EAS (J-L).

**Supplementary Figure 10 Population structure centers by country of birth:** UK Biobank continental ancestry group centers in the correspondence analysis, colored by K-means population clusters, overlapping with country of birth data in grey: AFR (A-C), EUR (D-F), SAS (G-I), EAS (J-L).

## Notes

### Competing Interest Statement

The authors have declared no competing interest.

