## Supplemental Figures 1-10 for "A framework for research into continental ancestry groups of the UK Biobank"

### Supplementary Figure 1: Continental ancestry PCA scree plots

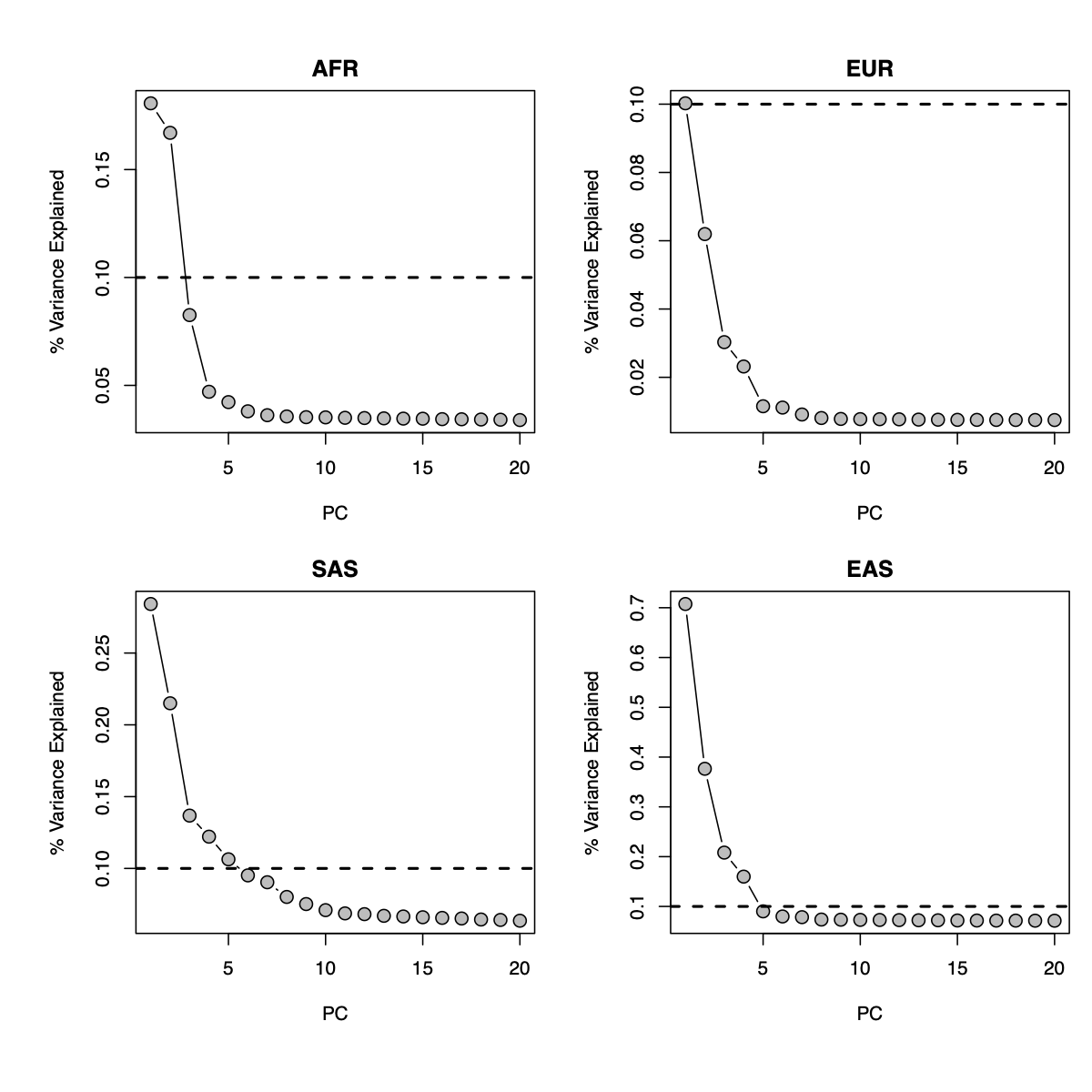

Legend: Scree plots illustrating the proportion of variation explained by each of the top 20 PCs, in each UKBB continental ancestry principal component analyses. The Scree plots were used to identify the number of top PCs to carry forward into the k-means clustering analysis. The continental ancestry supergroups are Africa (AFR), Europe (EUR), South Asia (SAS), and East Asia (EAS). The number of PCs selected as top PCs are AFR = 4, SAS = 5, EAS = 4, EUR = 5.

### Supplementary Figure 2: K-means k selection with silhouette analysis

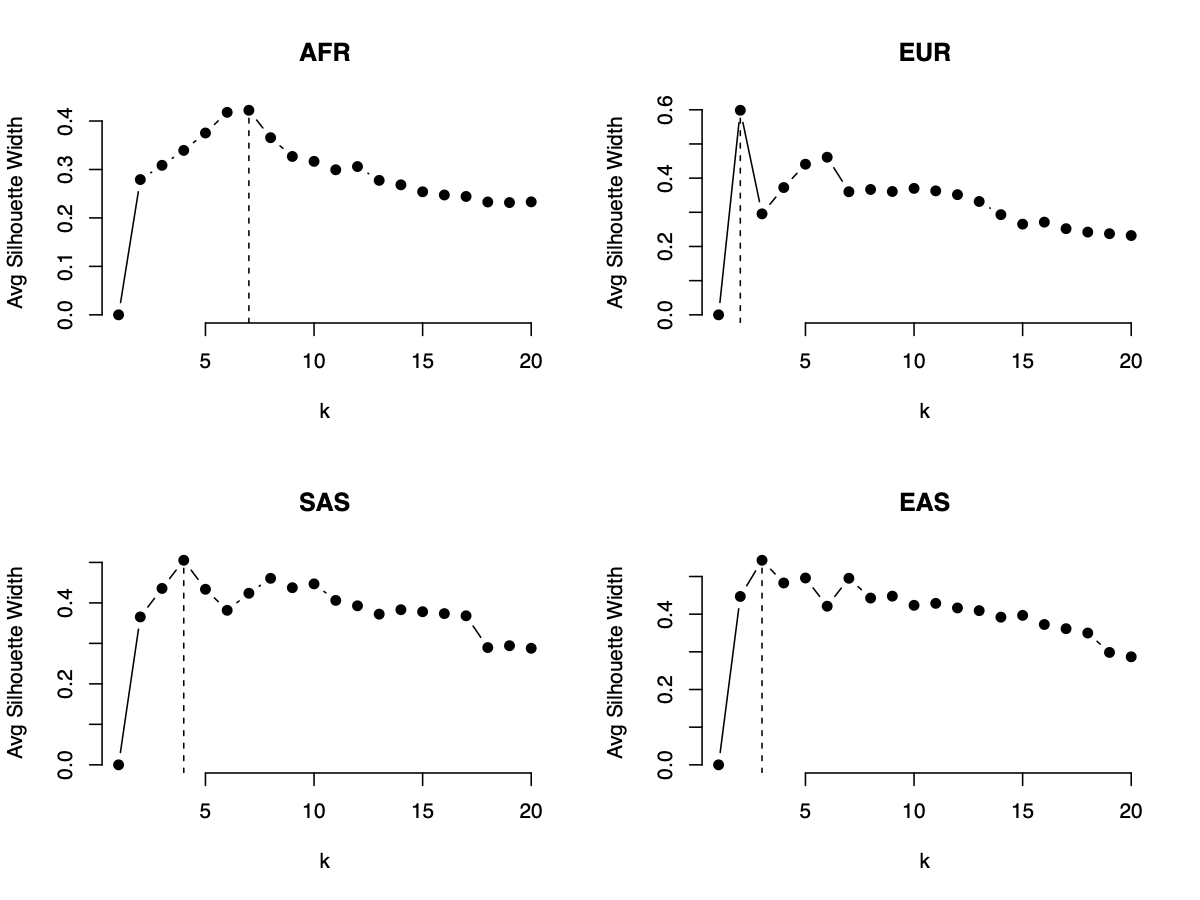

Legend: Selection of an optimum number of k clusters in the k-means analysis of the top PCs, by silhouette analysis. A silhouette plot for each UKBB continental supergroup (AFR) African, (EUR) European, (SAS) South Asian, and (EAS) East Asian is provided. The x-axis indicates the number of k clusters evaluated, and the y-axis provides an estimate of the average silhouette width (ASW). ASW is an estimation of cluster quality, or intra- and inter- cluster distances derived from a partitioning around medoids (PAM). The optimum number of k clusters in each UKBB continental supergroup were identified as AFR = 7, EUR = 2, SAS = 4, and EAS = 3.

### Supplementary Figure 3: UKBB continental ancestry group PCs with K-means clusters

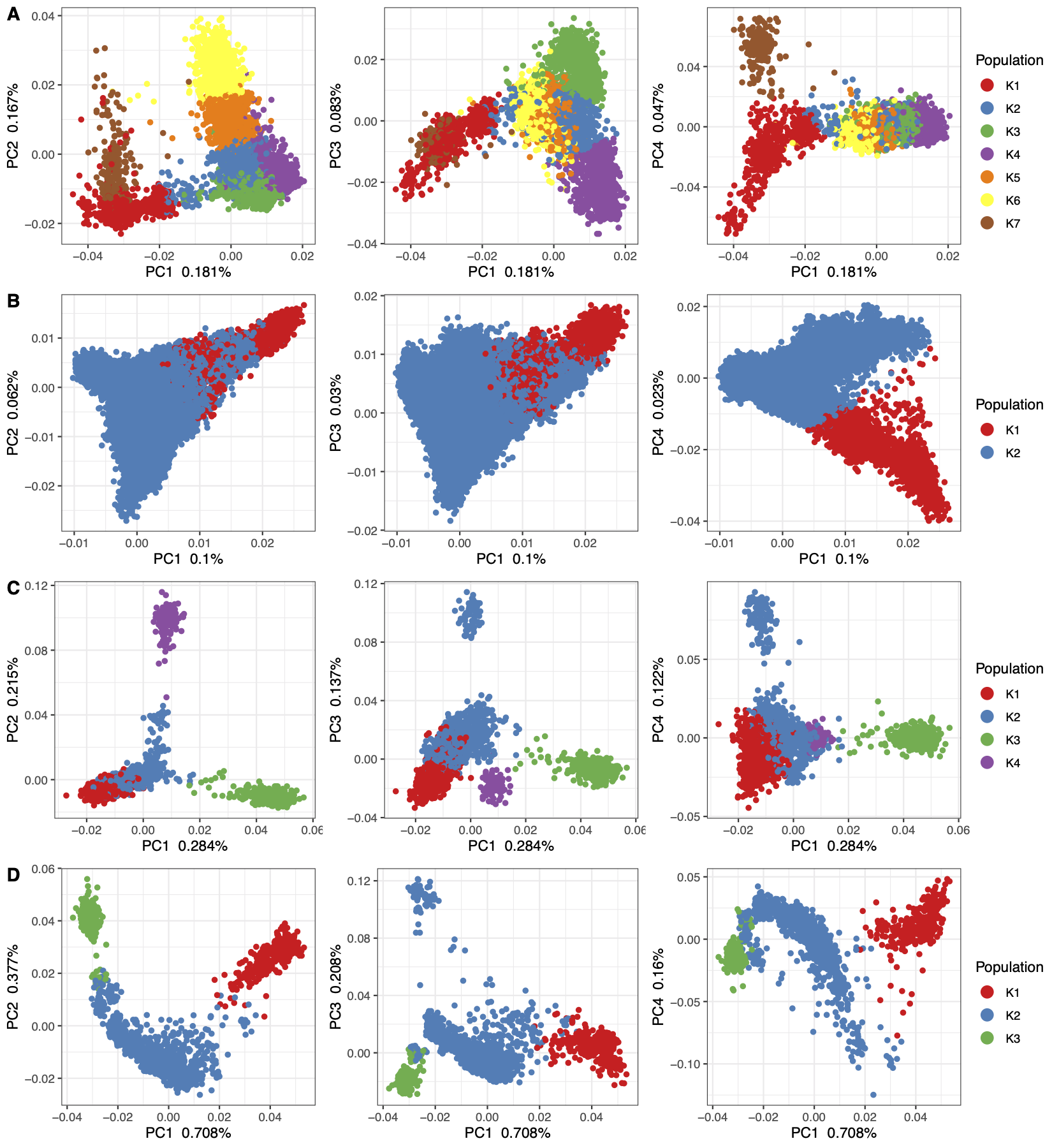

Legend: UK Biobank continental ancestry group PCs 1-4 with k-means groups colour coded: AFR (A), EUR (B), SAS (C), EAS (D).

### Supplementary Figure 4: Population structure by UN defined geographic region.

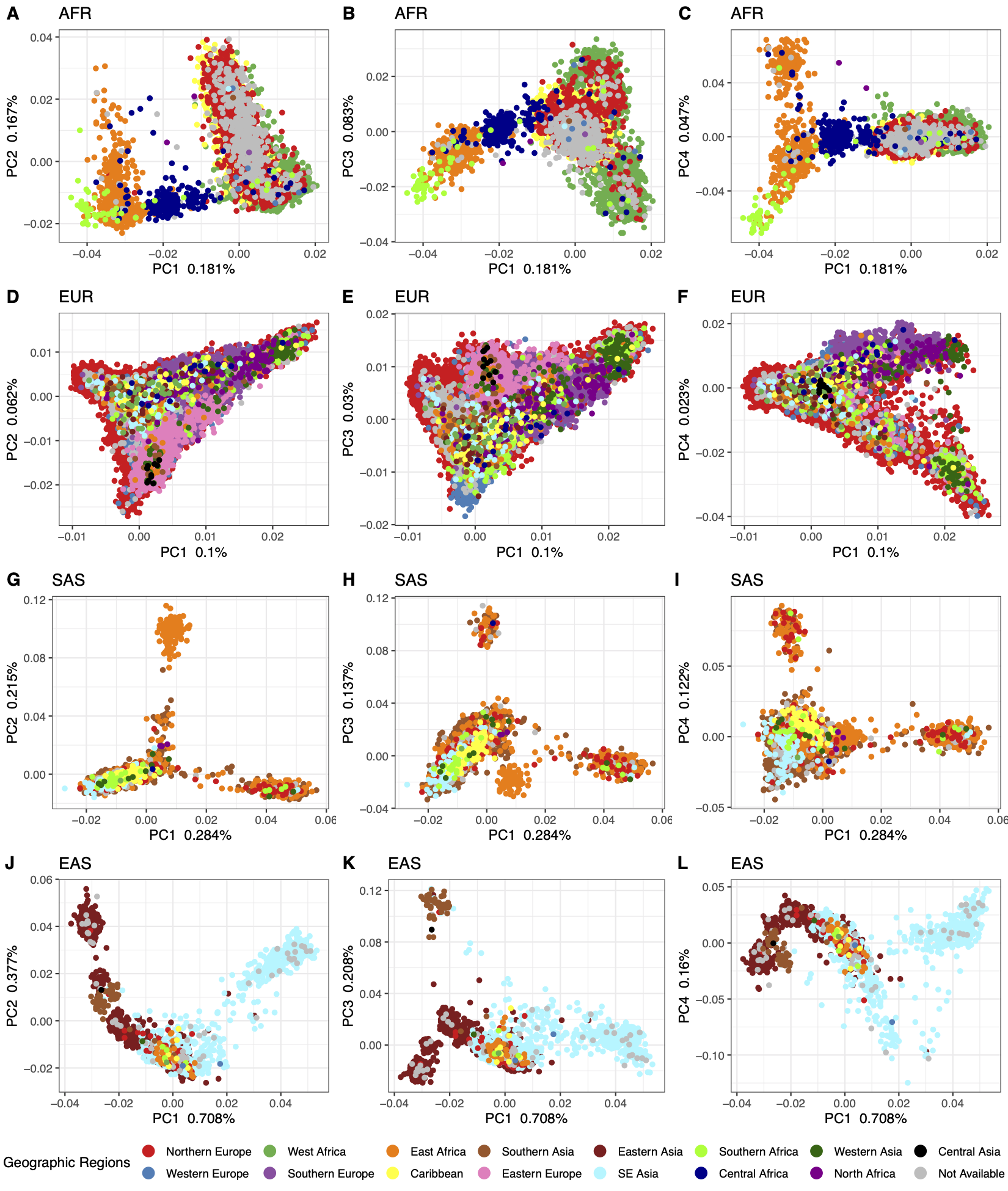

Legend: UK Biobank continental ancestry group PCs 1-4 with regions of birth colour coded: AFR (A), EUR (B), SAS (C), EAS (D).

### Supplementary Figure 5: Population structure centers, as defined by UN geographic region

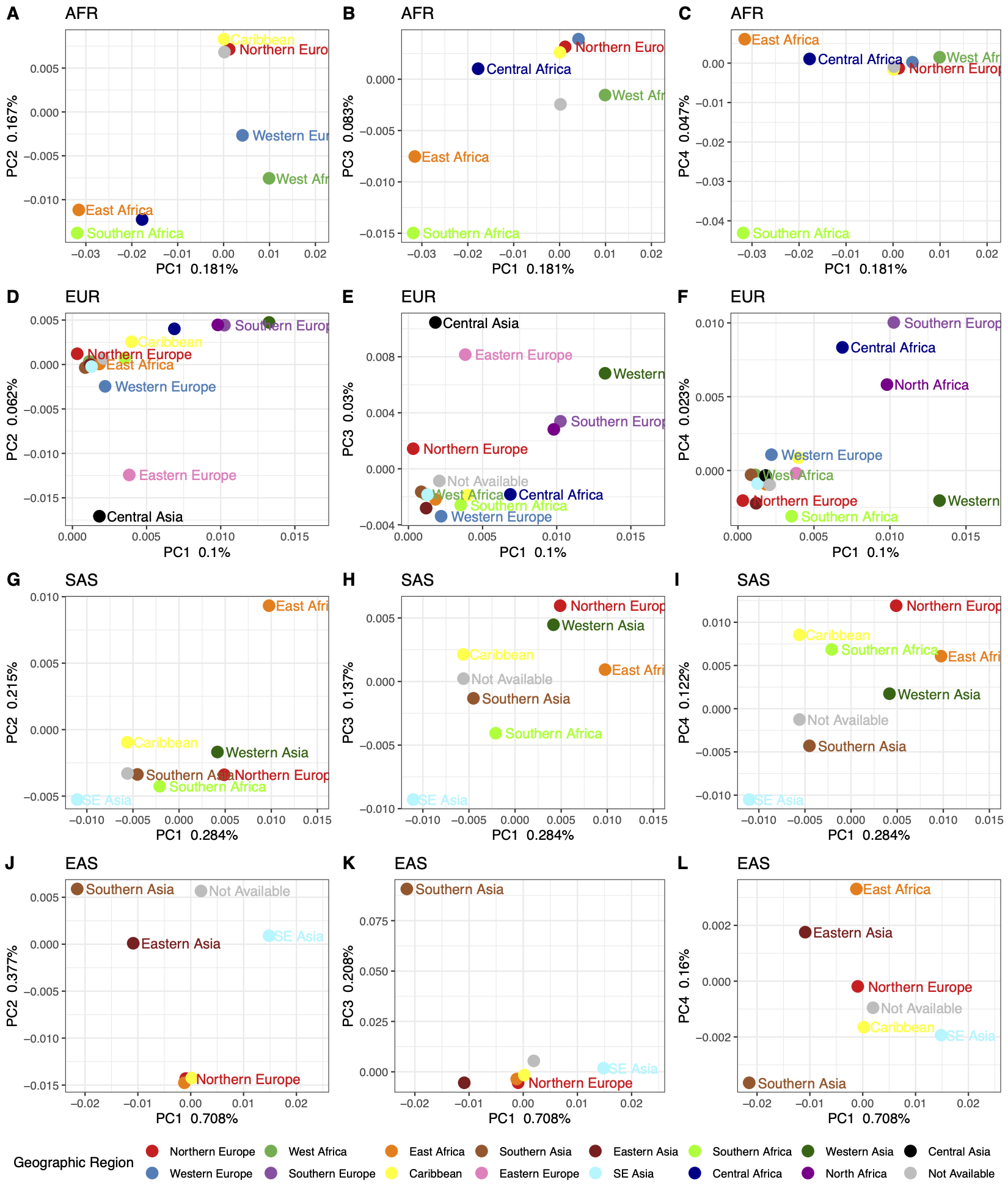

Legend: UK Biobank continental ancestry group PCs 1-4 with region of birth centers (averaged across all individuals from each ROB) colour coded: AFR (A-C), EUR (D-F), SAS (G-I), EAS (J-L).

### Supplementary Figure 6: Population structure by country of birth in AFR by region

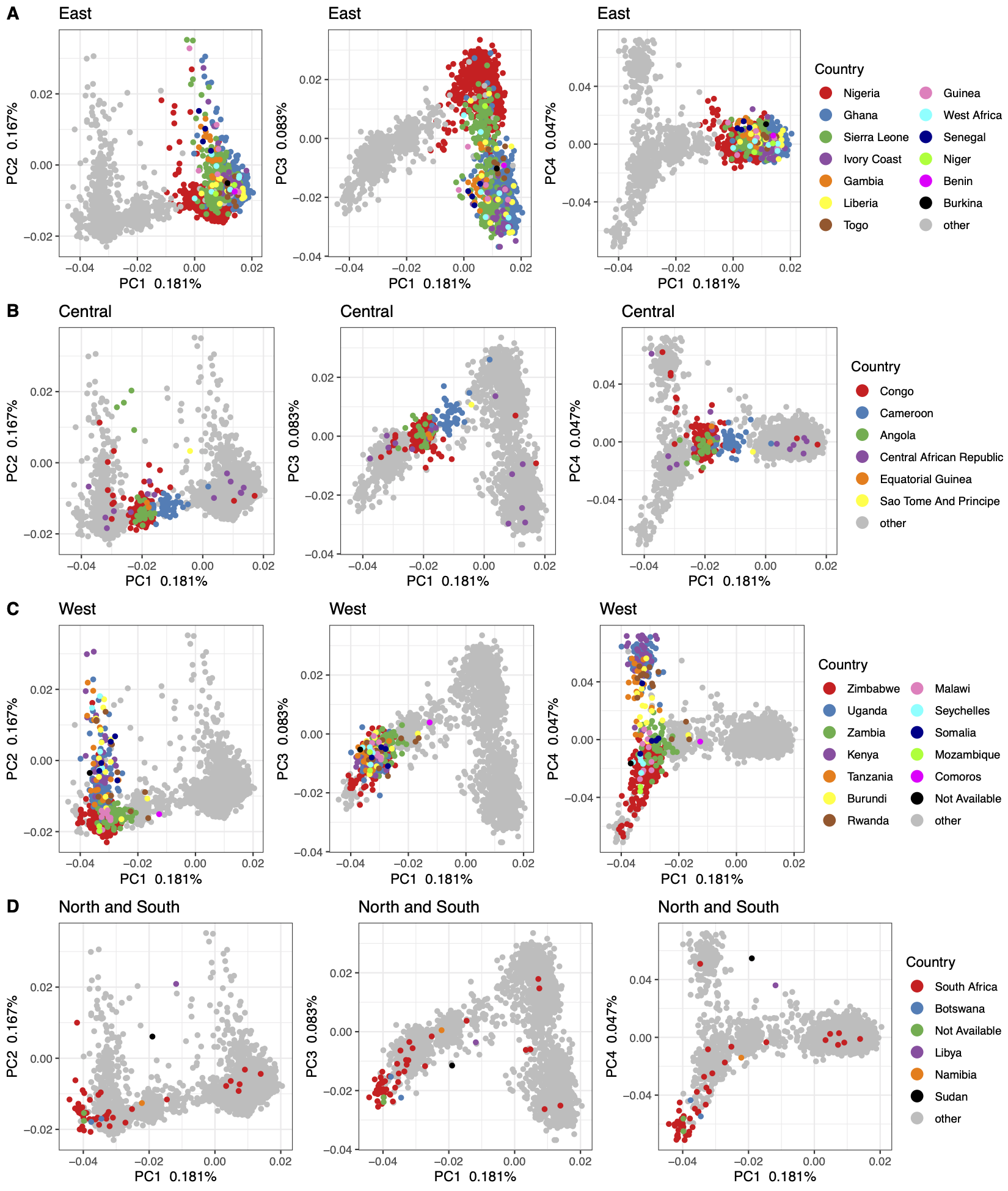

Legend: UK Biobank continental ancestry group PCs 1-4 for the AFR CAG divided by UN regions of birth: Eastern (A), Central (B), Western (C), Northern/Southern (D). Samples are colour coded by their country of birth.

### Supplementary Figure 7: Population structure by country of birth in EUR by region

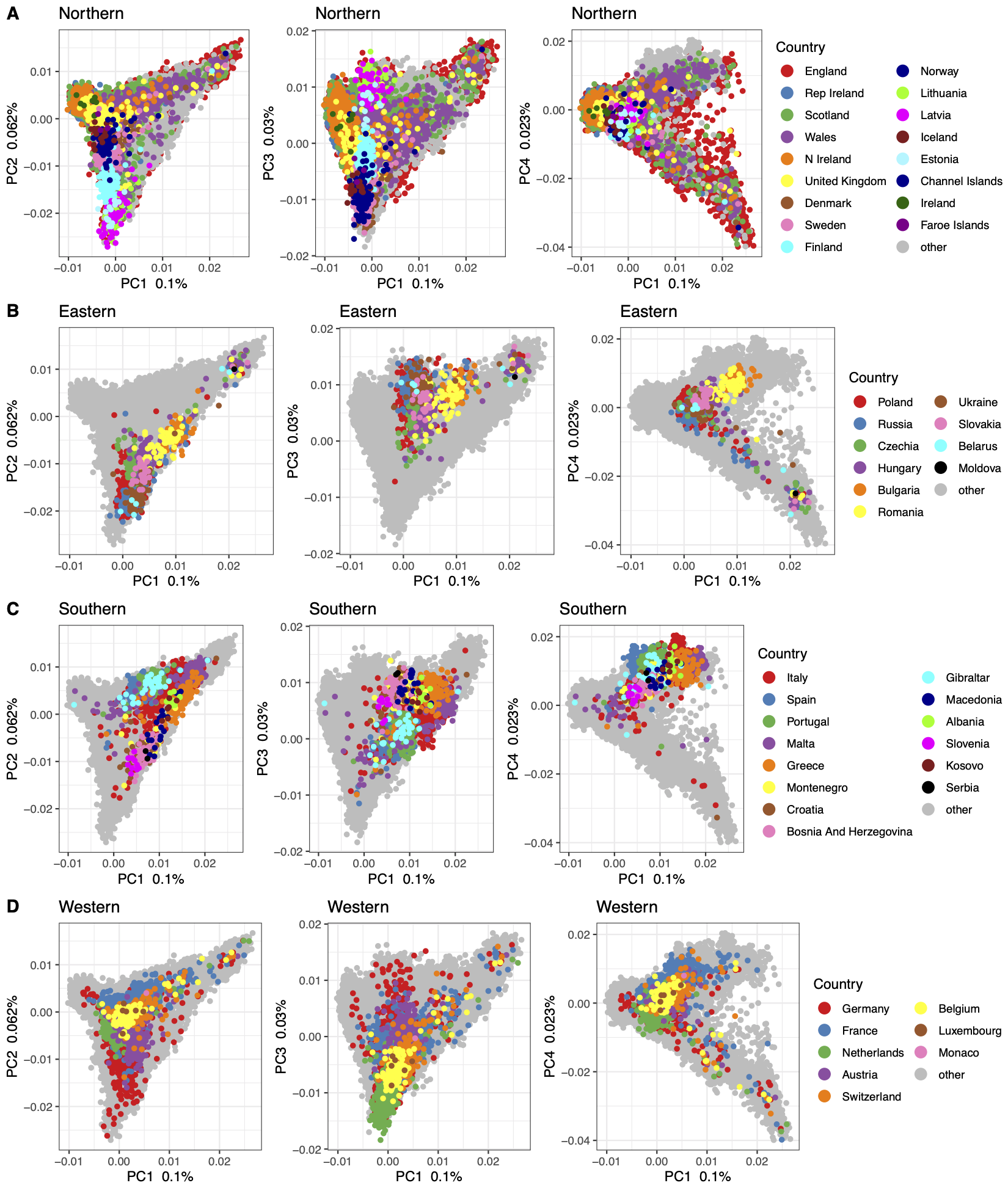

Legend: UK Biobank continental ancestry group PCs 1-4 for the EUR CAG divided by UN regions of birth: Northern (A), Eastern (B), Southern (C), Western (D). Samples are colour coded by their country of birth.

### Supplementary Figure 8: Population structure by country of birth in SAS and EAS

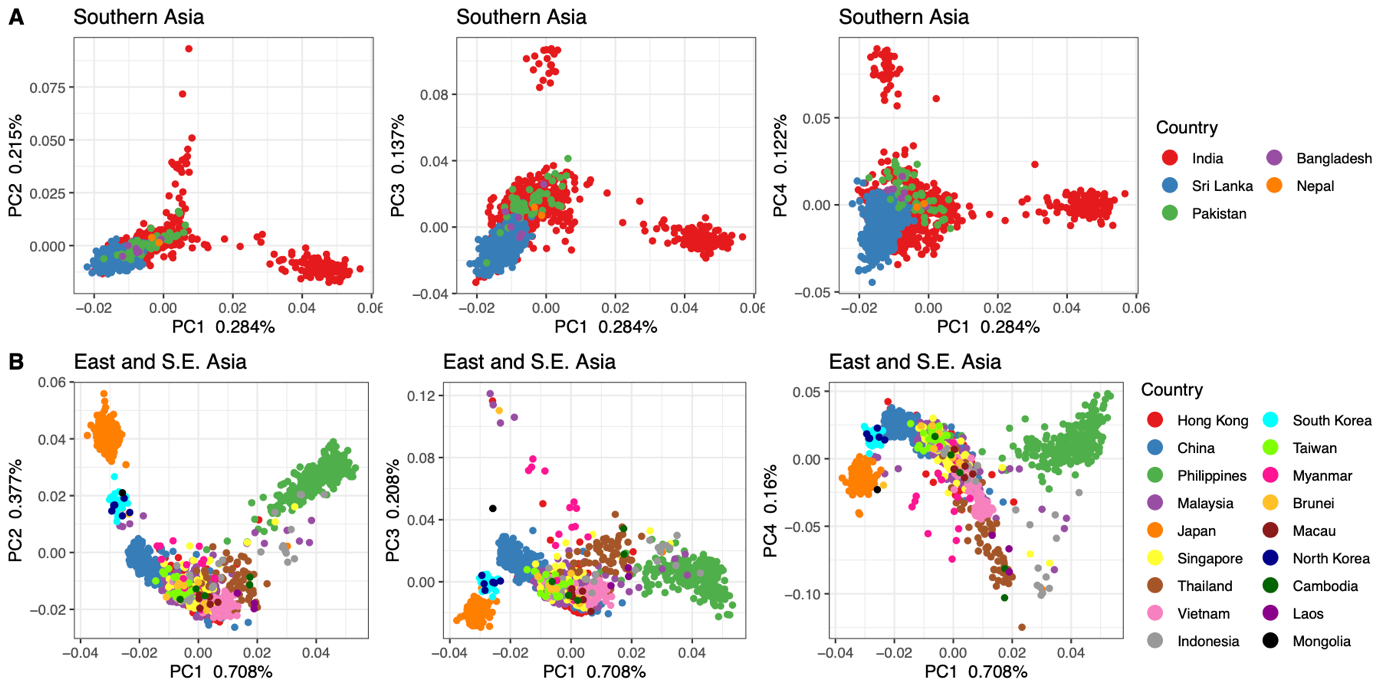

Legend: UK Biobank continental ancestry group PCs 1-4 for the SAS and EAS CAGs divided by UN regions of birth: Southern Asia (A), East Asia and South-East Asia (B). Samples are colour coded by their country of birth.

### Supplementary Figure 9: Population structure centers by country of birth

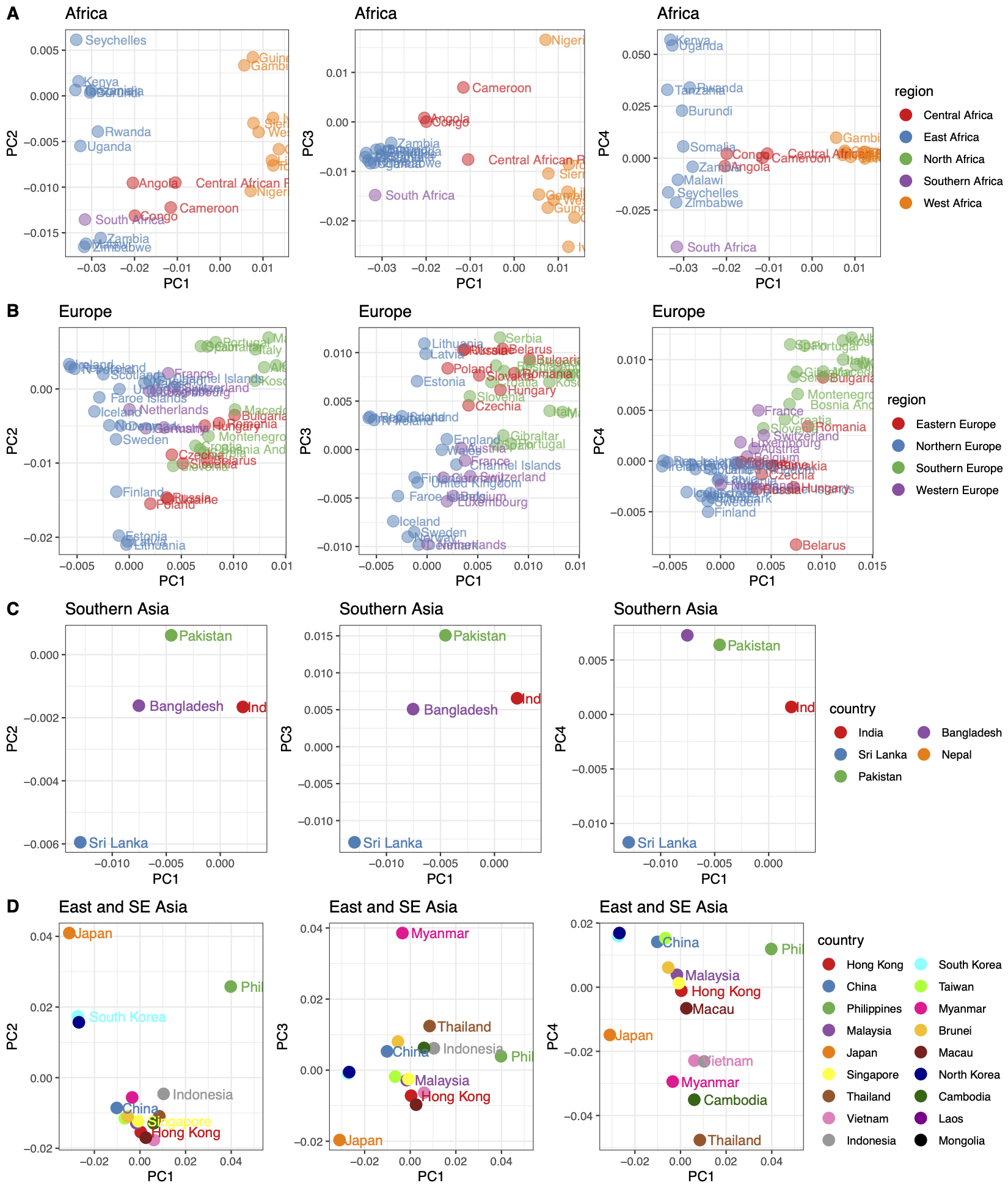

Legend: UK Biobank continental ancestry group centers colored by the country of birth: AFR (A-C), EUR (D-F), SAS (G-I), EAS (J-L).

### Supplementary Figure 10: Population structure centers by country of birth

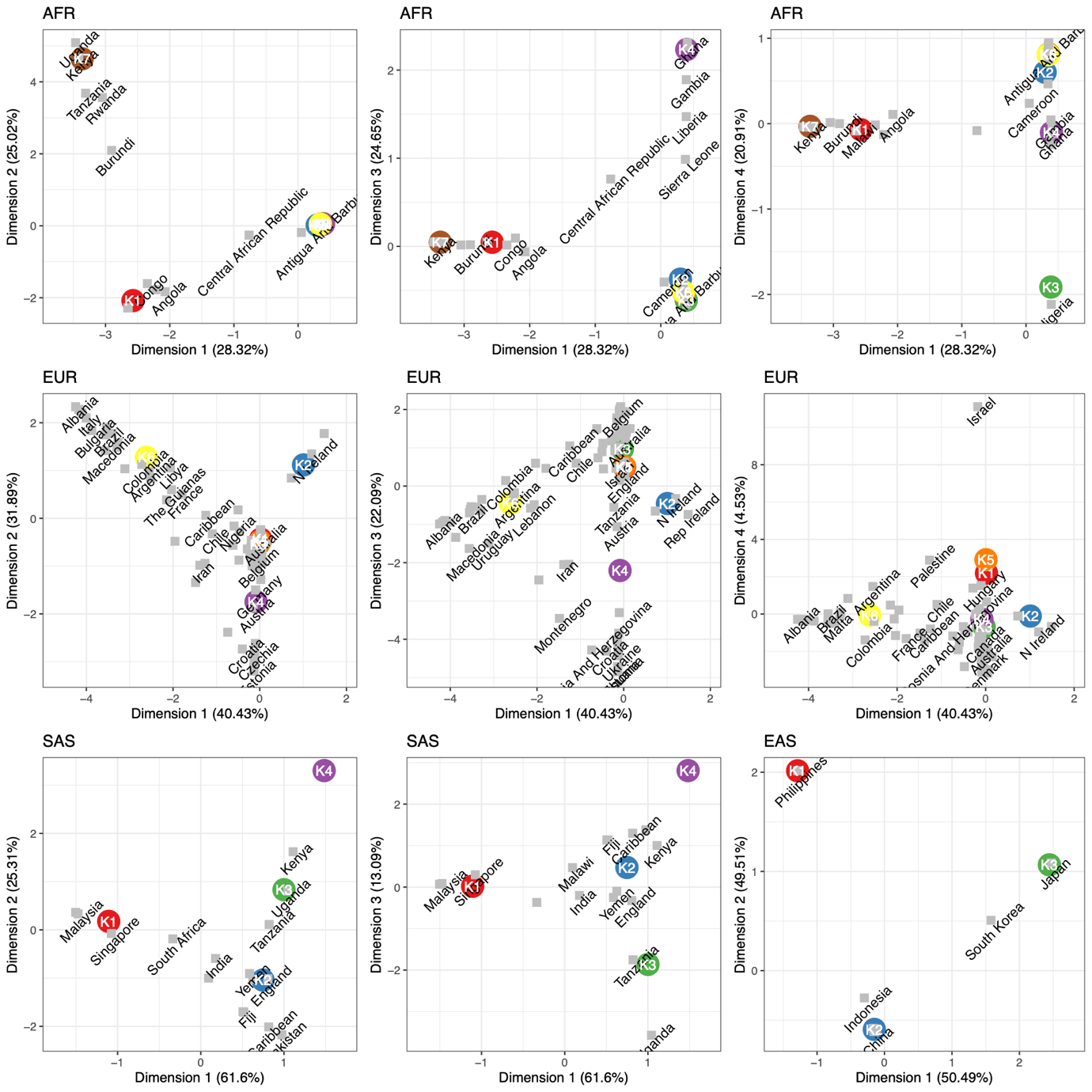
 Legend: UK Biobank continental ancestry group centers in the correspondence analysis, coloured by k-means groups, overlapping with country of birth data in grey: AFR (A-C), EUR (D-F), SAS (G-I), EAS (J-L).
